# Plasmid characterization in bacterial isolates of public health relevance in a tertiary healthcare facility in Kilimanjaro, Tanzania

**DOI:** 10.1101/2022.03.17.484711

**Authors:** Lameck Pashet Sengeruan, Marco van Zwetselaar, Happiness Kumburu, Frank M. Aarestrup, Katharina Kreppel, Elingarami Sauli, Tolbert Sonda

## Abstract

Plasmids are infectious double stranded DNA molecules that are found within bacteria. Horizontal gene transfer promotes successful spread of different types of plasmids within or among bacteria species, making their detection an important task for guiding clinical treatment. We used whole genome sequenced data to determine the prevalence of plasmid replicons in clinical bacterial isolates, the presence of resistance and virulence genes in plasmids, and the relationship between resistance and virulence genes within each plasmid. All bacterial sequences were de novo assembled using Unicycler before extraction of plasmids. Assembly graphs were submitted to Gplas+plasflow for plasmid prediction. The predicted plasmid components were validated using PlasmidFinder.

A total of 159 (56.2%) out of 283 bacterial isolates were found to carry plasmids, with *E. coli, K. pneumoniae and S. aureus* being the most prevalent plasmid carriers. A total of 27 (87.1%) combined plasmids were found to carry both resistance and virulence genes compared to 4 (12.9%) single plasmids. No statistically significant correlation was found between the number of antimicrobial resistance and virulence genes in plasmids (r =-0.25, p > 0.05). Our findings show a relatively high proportion of plasmid-carrying isolates suggesting selection pressure due to antibiotic use in the hospital. Co-occurrence of antibiotic resistance and virulence genes in clinical isolates is a public health relevant problem needing attention.

## Introduction

Plasmids are circular, double-stranded DNA molecules that occur naturally in bacterial cells [1], whose genes often provide evolutionary advantages for bacteria, such as antimicrobial resistance and/or virulence [2,3]. Plasmids are important vehicles in disseminating and acquiring antibiotic resistance and virulence, and can thus constitute a major burden on human health [4]. Recent studies have suggested that the prevalence of antimicrobial resistance (AMR) is higher in Low- and Middle-income Countries (LMICs) compared to European countries and the United States [5,6]. There is however, limited knowledge regarding the dissemination of antibiotic resistance genes (ARGs) and virulence among clinical isolates in Sub-Saharan Africa (SSA). This study was conducted to determine the proportion of bacterial isolates carrying plasmids, to identify plasmids that mediate resistance and virulence genes, and to investigate the relationship between antimicrobial resistance genes and virulence genes within plasmids using whole genome sequence data from bacterial isolates among inpatients at Kilimanjaro Christian Medical Centre (KCMC) in Tanzania.

## Materials and methods

### Study setting, Whole-genome sequencing and library preparation

Kilimanjaro Christian Medical Centre (KCMC) is one of Tanzania’s five zonal referral hospitals, located in Moshi, northern Tanzania. KCMC has a bed capacity of 650 and serves a catchment area of about 15 million people. It serves around 500 outpatients daily, from different parts of Tanzania [7]. The whole genome sequence data that was analyzed originated from a prospective cross- sectional study that was conducted at KCMC between August 2013 and August 2015. In this study, a total of 56 stool, 122 sputum, 126 blood and 286 wound swabs (wound/pus) clinical samples, with patients’ clinical and socio-demographic characteristics, were collected from 575 patients admitted to KCMC hospital [8,9]. A written informed consent was obtained from each participant and from parents or guardians of children before enrolled into the study.

Collected specimens were taken to the microbiology unit at Kilimanjaro Clinical Research Institute (KCRI) for culture and identification of bacterial isolates. Out of 590 specimens collected, 249 were culture positive, resulting in 377 isolates [8]. All bacterial isolates were sequenced in the KCRI genomics lab, and all sequences were archived on the KCRI compute cluster. In brief, whole genome sequencing (WGS) was performed for genomic DNA that was extracted from cultures of bacterial isolates using the Easy-DNA Extraction Kit (Invitrogen®). Short-read WGS was performed using the Illumina MiSeq platform (Illumina Inc.). Libraries for Illumina sequencing were constructed using the Illumina Nextera XT kit (Illumina Ltd., San Diego, CA, USA) according to the manufacturer’s recommendations. The libraries were sequenced on Illumina MiSeq platform using the 2 × 250bp paired-end protocol, previously reported by Kumburu et al [8] & Sonda et al [9]. For the purpose of this study, a total of 283 bacterial whole genomes isolates with associated metadata were retrieved for analysis. Additional ethical approval was obtained from the Ifakara Health Institute Research Ethics Committee (IHI/IRB/No: 14-2021) for plasmid characterization.

### Bioinformatics Analyses

#### Quality Control and Trimming of Illumina Sequences

The following steps were followed: (i) All bacterial raw reads were submitted to in-house bacterial analysis pipeline (BAP), available at https://github.com/zwets/kcri-cge-bap. Assembly was performed using SKESA 2.4.0 [10]. (ii) All resulting assemblies were then processed in batch by the Genome Taxonomy Database Toolkit (GTDB-Tk) 0.3.2 [11] for detailed taxonomic assignment. (iii) Metrics produced by the BAP and GTDB-Tk were then used to assess the quality of each assembly. Assessment was based on read counts, coverage depth, assembly structure (contig count, N1, N50, L50), deviation of assembly length from reference, GTDB alignment fraction, and GTDB Multi-Locus Sequence Alignment (MSA) coverage. A six-point scale was used for assembly quality rating: 0 (Unusable), 1 (Mix), 2 (Bad), 3 (Usable), 4 (Good), and 5 (Excellent). (iv) Finally, categories 0 to 2 were excluded, while categories 3 through 5 were used for subsequent analysis. Every assembly in these categories was for a single isolate that had (nearly) complete genome coverage, at sufficient sequencing depth.

#### Plasmid extraction and validation

Raw reads assembly was repeated with Unicycler 0.4.7 [12] for its ability as a “SPAdes optimiser” to produce long and, in the ideal case, circular contigs. Assembly graphs (GFA) were submitted to Gplas+plasflow for plasmid prediction. Gplas 0.6.1 [13] + Plasflow 1.1 [14] take into account the connected components in the assembly graph when predicting plasmids. The components predicted to be plasmids were extracted from the assemblies and submitted to PlasmidFinder version 1.3 [15] for validation.

#### Identification of Plasmid-Mediated Antibiotic Resistance Genes (ARGs) and Virulence genes

To identify antibiotic resistance and virulence genes carried in plasmids, the assembled putative plasmid sequences for each isolate were submitted to Resfinder 4.0 [16] and VirulenceFinder 1.4[17] respectively. In both Resfinder and VirulenceFinder, 90% identity and 60% coverage settings to call a gene were selected.

#### Statistical analysis

Stata 14 (College Station, TX, 77845, USA) was used for descriptive statistics and determination of the relationship between antimicrobial resistance and virulence genes in plasmids.

## Results

### Study population

In total, 128 patients whose whole genome bacterial isolates were analyzed were included in this study (Table 1). One-hundred twenty eight patients were plasmid positive isolates. The mean age in years (SD) was 46.2 (18.0). Male patients were 77 (60.2%), females were 47 (36.7%) and 4 (3.1%) missed gender identification. A total of 62 (48.4%) patients were admitted to surgical ward, 9 (7.1%) surgical ICU, 52 (40.6%) medical ward, 4 (3.1%) medical ICU ward, and 1 (0.8%) missed ward admission identification. Eighty-seven (67.9%) specimens were swabs, 19 (14.8%) were stool, 13 (10.2%) were sputum, 8 (6.3%) were blood and 1(0.8%) specimen missed identification. A total of 28 (21.9%) patients were diabetic, 6 (4.7%) were cancer patients, 6 (4.7%) were suffering from TB, 2 (1.5%) were HIV positive and 86 (67.2%) were others: Of those 61 (47.7%) had no underlying conditions and 25 (19.5%) had other underlying conditions. Of the wound swabs, twenty-four (18.8%) were from patients with diabetic wounds, 11 (8.6%) were from burn wounds, 10 (7.8%) were from post-surgical wounds, 6 (4.6%) were from motor traffic accidents wounds and 77 (60.2%) were others: Of those 35 (27.3%) other wounds and 42 (32.8%) had no wounds.

**Table 1.** Demographic and clinical characteristics of patients that samples were taken from

| Patient characteristics | Missing <sup>a</sup> | Total (%) |
| --- | --- | --- |
| Number of patients |  | 128 (100) |
| Mean age in years (SD) |  | 46.2 (18.0) |
| Gender | 4 (3.1) |  |
| Female |  | 47 (36.7) |
| Male |  | 77 (60.2) |
| Ward of admission | 1 (0.8) |  |
| Surgical |  | 62 (48.4) |
| Surgical ICU |  | 9 (7.1) |
| Medical |  | 52 (40.6) |
| Medical ICU |  | 4 (3.1) |
| Specimen collected | 1 (0.8) |  |
| Blood |  | 8 (6.3) |
| Sputum |  | 13 (10.2) |
| Stool |  | 19 (14.8) |
| Swab |  | 87 (67.9) |
| Underlying conditions |  |  |
| Cancer |  | 6 (4.7) |
| Diabetes |  | 28 (21.9) |
| HIV |  | 2 (1.5) |
| TB |  | 6 (4.7) |
| Others |  | 86 (67.2) |
| Type of wound |  |  |
| Burn wound |  | 11 (8.6) |
| Diabetic wound |  | 24 (18.8) |
| Motor traffic wound |  | 6 (4.6 ) |
| Post-surgical wound |  | 10 (7.8) |
| Others |  | 77 ( 60.2) |
| History of hospitalization | 8 (6.3) |  |
| No |  | 78 (60.9) |
| Yes |  | 42 (32.8) |
| Patient hospital transfer | 3 (2.3) |  |
| No |  | 44 (34.4) |
| Yes |  | 81 (63.3) |
| Patient ward transfer | 5 (3.9) |  |
| No |  | 110 (85.9) |
| Yes |  | 13 (10.2) |
| Median time in days stayed in the hospital before survey (IQR) | 8 (6.3) | 8 (4-11.5) |

SD, standard deviation; ICU, intensive care unit; IQR, interquartile range; Missing <sup>a</sup>, were the missing values in each variable.

Seventy-eight (60.9%) patients had no history of hospitalization, 42 (32.8%) had hospitalization history and 8 (6.3%) missed identification. A total of 81 (63.3%) patients were transferred from another hospital, 44 (34.34%) were not and 3 (2.3%) missed identification. Among all participants, the median number of days stayed in the hospital before the survey was 8 days (Table 1).

### Proportion of bacterial species carrying plasmids

A total of 283 whole genome bacterial sequences were analyzed. One hundred fifty-nine (56.2%) bacterial isolates were detected to carry plasmids. Out of 159 plasmids, 94 non-repetitive plasmids were predicted. Of 94 plasmids, 48 (51.1%) were single plasmids and 46 (48.9%) were combined plasmids (two or more recombined plasmids). *K. pneumoniae* isolates were the most carriers of combined plasmids (17, 28.3%), followed by *S. aureus* (15, 25.0%) and *E*.*coli* (15, 25.0%). *E*.*coli* isolates were the most single plasmid carriers (23, 23.2%), followed by *S. aureus* (15, 15.2%) and *P. mirabilis* (14,14.4%) (Table 2).

**Table 2.** Proportion of bacterial isolates carrying plasmids

|  |  |  | Plasmids |  |
| --- | --- | --- | --- | --- |
|  |  |  | Single | Combined <sup>b</sup> |
| Species | N | % | N | N |
| *Enterobacter asburiae* | 1 | 0.6 | 1 | 0 |
| *Enterobacter cloacae* | 3 | 1.9 | 1 | 2 |
| *Enterobacter hormaechei* | 10 | 6.3 | 6 | 4 |
| *Enterobacter kobei* | 1 | 0.6 | 0 | 1 |
| *Enterobacter roggenkampii* | 1 | 0.6 | 1 | 0 |
| *Enterobacter soli* | 1 | 0.6 | 1 | 0 |
| *Enterobacter sp. n18-03635* | 1 | 0.6 | 1 | 0 |
| *Enterococcus faecalis* | 7 | 4.4 | 5 | 2 |
| *Enterococcus faecium* | 3 | 1.9 | 2 | 1 |
| *Enterococcus gallinarum* | 1 | 0.6 | 1 | 0 |
| *Escherichia coli* | 38 | 23.9 | 23 | 15 |
| *Klebsiella Michiganensis* | 2 | 1.3 | 1 | 1 |
| *Klebsiella oxytoca* | 2 | 1.3 | 1 | 1 |
| *Klebsiella pneumoniae* | 25 | 15.7 | 8 | 17 |
| *Klebsiella variicola* | 2 | 1.3 | 2 | 0 |
| *Micrococcus sp. Kbs0714* | 1 | 0.6 | 1 | 0 |
| *Morganella morganii* | 4 | 2.5 | 4 | 0 |
| *Proteus columbae* | 1 | 0.6 | 1 | 0 |
| *Proteus mirabilis* | 14 | 8.8 | 14 | 0 |
| *Proteus penneri* | 1 | 0.6 | 1 | 0 |
| *Proteus vulgaris* | 1 | 0.6 | 1 | 0 |
| *Pseudomonas aeruginosa* | 2 | 1.3 | 2 | 0 |
| *Shewanella algae* | 1 | 0.6 | 1 | 0 |
| *Staphylococcus aureus* | 30 | 18.9 | 15 | 15 |
| *Staphylococcus capitis* | 1 | 0.6 | 1 | 0 |
| *Staphylococcus epidermidis* | 1 | 0.6 | 1 | 0 |
| *Staphylococcus haemolyticus* | 3 | 1.9 | 3 | 0 |
| *Staphylococcus hominis* | 1 | 0.6 | 0 | 1 |

N, total number of isolates/plasmids; combined <sup>b</sup>, two or more plasmids merged or fused together.

### Plasmids concurrently mediating resistance and virulence genes

A total of 31 plasmids were identified to carry both resistance and virulence genes, of which 27 (87.1%) were combined plasmids and 4 (12.9%) were single plasmids (Tables 3 and 4). All four single plasmids were carried by *E. coli*. Resistance gene *Sul1* was found the most common across three single plasmids IncFII, IncQ1 and IncFII(pRSB107). Virulence genes iucC *and iutA* were also seen the most common across three single plasmids such as IncQ1, IncFII(pRSB107) and IncFIA (Table 3).

**Table 3.** Single plasmids mediating both resistance and virulence genes

| Single plasmids | Resistance genes | Virulence genes |
| --- | --- | --- |
| IncFII | *aac(3)-IIa,aadA5,blaCTX-M-15,dfrA17,qacE,sul1,tet(B)* | *traT* |
| IncQ1 | *aph(3'')-Ib,aph(6)-Id,blaTEM-1B,dfrA7,qacE,sul1,sul2* | *cea,focCsfaE,focG,focI,iha,ireA,iucC,iutA,mchB,mchC,mchF,mcmA,papA_F48,papC,sat* |
| IncFIA | *sitABCD,tet(A)* | *iucC,iutA,sitA* |
| IncFII(pRSB107) | *dfrA5,qacE,sul1,sul2* | *capU,iroN,iss,iucC,iutA,mchB,mchC,mchF,mcmA,vat* |

**Table 4.**
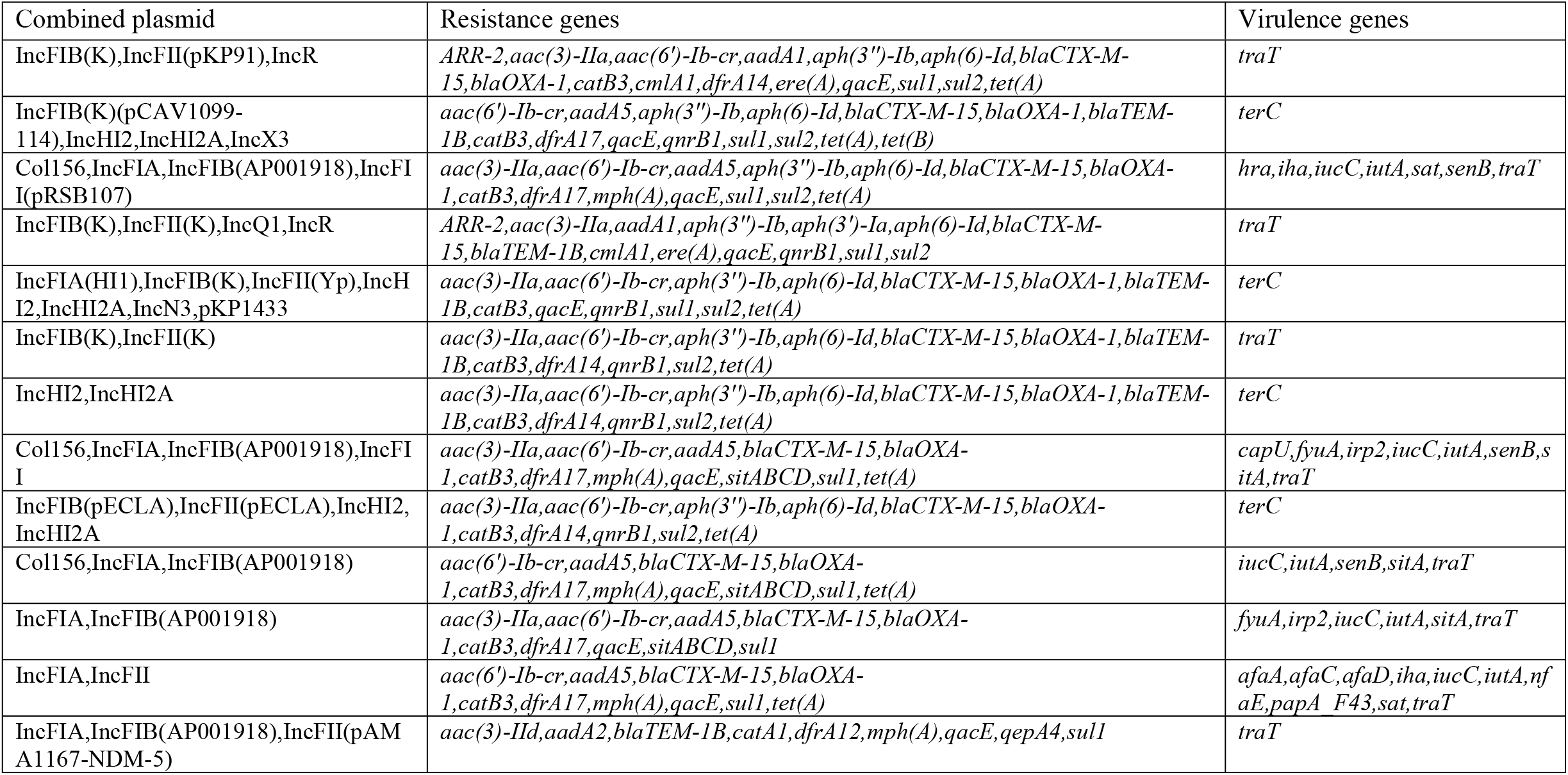
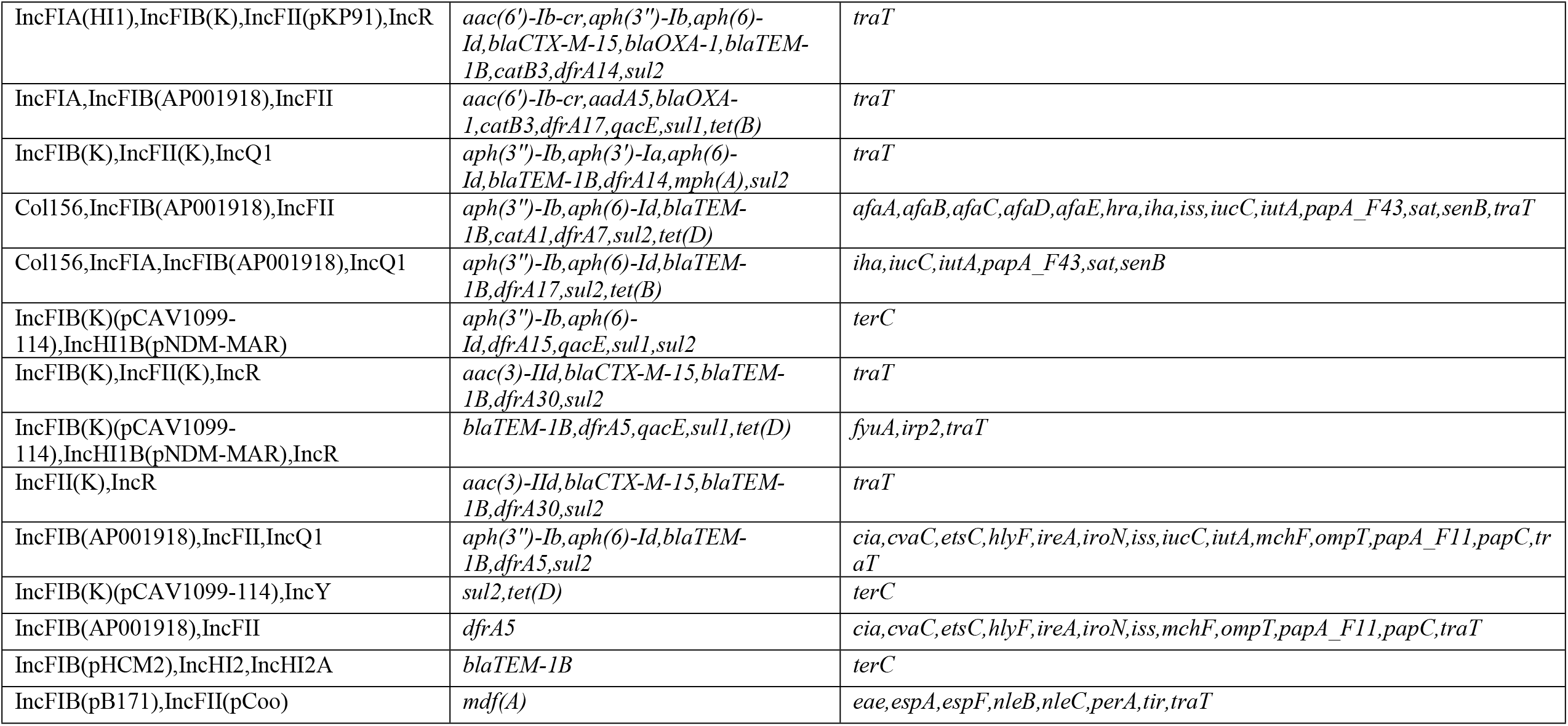
Combined plasmids mediating both resistance and virulence genes

Among the 27 combined plasmids, 12 (44.4%) were carried by *E*.*coli* isolates, 10 (37.1%) by *K. pneumoniae* isolates, 2 (7.4%) by *E. hormaechei* isolates, 1 (3.7%) by *E. cloacae* isolate, 1(3.7%) by *K. oxytoca* isolate and 1 (3.7%) by *K. michiganensis* isolate. Virulence gene *traT* was seen in 19 (70.4%) of the 27 combined plasmids, followed by *terC* which was identified in 7 (25.9%) plasmids. Resistance genes in combined plasmids, *sul2* was observed in 17 (62.9%) plasmids, followed by *blaTEM-1B* in 15 (55.6%) plasmids, followed by *blaCTX-M-15* in 14 (51.9%) plasmids and *blaOXA-1* in 13 (48.1%) plasmids (Table 4).

### Correlation between antimicrobial resistance and virulence genes

We explored the relationship between the number of antibiotic resistance genes and virulence genes in 27 combined plasmids using Pearson correlation. There was an inconclusive negative relationship between antibiotic resistance and virulence genes existence in plasmids (r =-0.25, p > 0.05).

## Discussion

In the present study a high proportion of clinical bacterial isolates from inpatients at KCMC hospital was found to carry plasmids. The present findings are in concordance with previous studies elsewhere [18]. The observed high carriage of plasmids by the analyzed isolates might plausibly be a reflection of resistance selection pressure due to high antibiotic exposure in hospital settings [19].

*E. coli* isolates were the most prevalent carriers of single plasmids followed by *S. aureus* and *P. mirabilis*. On other hand, *K. pneumonia* were the most prevalent carriers of combined plasmids, followed by *S. aureus* and *E. coli*. The present study findings are in line with a study results in a tertiary care hospital in south India [18]. Possible explanation could be that the mentioned bacterial species have great medical relevance and thus are relatively highly isolated in hospital settings compared to other species [20,21]. However, the present study findings show a larger proportion of *P. mirabilis* carrying plasmids than the study in south India. This difference might be due to the fact that majority of the present study isolates were from wound specimens in which *P. mirabilis* were identified [22,23].

This study identified bacterial species with low plasmid prevalence including *Enterobacter sp. n18- 03635, Enterobacter kobei, Klebsiella variicola* and *Klebsiella oxytoca*. The study findings are consistent with other studies conducted in Canada, Greece and Mexico. The study findings are consistent with other studies conducted in Canada, Greece and Mexico [24–26]. Interestingly, the study observed other low plasmid prevalence species that were reported elsewhere in soil samples, fish flesh samples [27–29] and pigeon flesh specimens [30] such as *Micrococcus sp. Kbs0714, Enterobacter soli* and *Proteus columbae*. The observed species with low plasmid prevalence might be due to rarity and in most cases misidentification [31,32]. However, reports of bacterial species with low plasmid indicates the possible emerging and transmission of bacterial pathogens in humans both in community and hospital settings [33].

Contrary to previous studies reporting IncF plasmid group in *E*.*coli* to carry resistance and virulence genes often [34] the present study, however shows IncQ1 carried the highest number of both resistance and virulence genes in *E. coli*, but the finding is in line with study conducted in Brazil [35,36]. This is possibly due to the fact that IncQ1 plasmids have high-level mobility, stability, replication at high copy number and transferred in wide range of bacterial species through conjugative plasmids [37–40].

In this study it was also identified that there are different combined plasmids ranging from two to seven plasmids. This is probably an indicative of bacterial evolution to adapt and thrive in hospitals where they are excessively exposed to antimicrobials, antiseptics and disinfectants [41–43]. A similar distribution of some combined plasmids in other regions carrying similar or different antibiotic resistance and virulence genes was noted in the present study. This suggests resistant bacteria arising in one geographical area can spread countrywide/worldwide either by direct exposure or through the food chain or climate change and the environment [6].

There was no significant relationship found in the present study between numbers of antibiotic resistance and virulence genes in plasmids (r = −0.25, p > 0.05), indicating acquisition of antibiotic resistance can induce the loss of virulence factors. Previous studies support this study finding [44], but does not agree with a study by Dionisio [45]. This discordance might be due to the fact that in other studies the relationship between resistance and virulence genes was determine at species level and were from gut and environmental samples [34].

## Limitations

We acknowledge there are a number of limitations in the present study that warrant careful interpretation. Bioinformatics analysis was performed on Illumina short reads, which limited the ability to assemble completed plasmid genomes, and consequently the ability to ‘tease out’ individual plasmids from assembled contigs. Assembly graphs were classified by Gplas+PlasFlow for plasmid prediction. As for any machine learning-based approach or indeed any method based on inference from similarity with known sequences, including tools such as PlasmidFinder, the predictive ability of the model is strongly dependent on the data in its reference database or training set. A bias toward plasmids in well-studied organisms is therefore likely.

## Conclusion

There is a high proportion of isolates carrying resistance and virulence plasmids. This shows a significant concern of AMR development and spread in Tanzanian health settings and other LMIC settings. With limited resources and health service capacities, the increasing AMR trends are expected to high impact on bacterial-associated mortalities and morbidities.

## Acknowledgements

The parent study whose data we analyzed in this study was funded by DANIDA with grant number DFC No. 12-007DTU. We would like to express our sincere gratitude to study participants whose specimens yielded the bacterial isolates analyzed.

## Authors’ contributions

LP, MvZ, HK, FMA and TS conceived the initial idea. LP, TS, KK and ES developed the study protocol. TS, MvZ and LP analyzed the data. LP wrote the first draft of the manuscript. TS, MvZ, KK, ES, HK and FM reviewed and edited drafts of the manuscript. LP, TS, MvZ, KK and ES, HK and FMA wrote the final Manuscript. All authors read and approved the final manuscript.

## Availability of data

All relevant data are within the manuscript and its Supporting Information files.

## Consent for publication

Not applicable

